# From Data Curation to Risk Reporting: A Pipeline for Polygenic Risk Scores

**DOI:** 10.64898/2026.08.12.743994

**Authors:** Pedro Victor Barbosa Araújo, Tayná da Silva Fiúza, José Eduardo Kroll, Rodrigo Lima Andrade, Daniel Henrique Ferreira Gomes, Leonardo Varuzza, Gustavo Antonio de Souza, Sandro José de Souza

## Abstract

Polygenic risk scores (PRS) have emerged as a powerful tool for quantifying genetic susceptibility to complex traits and diseases. However, their calculation and interpretation require standardized data curation, robust statistical methods, and clear reporting strategies. In this work, we present an integrated pipeline designed to address these challenges. The pipeline begins with the construction of a curated genotype/phenotype database derived from public repositories, ensuring that only phenotypes with appropriate metadata, statistical distributions, and ethical suitability are retained. The final dataset comprises 2,346 phenotypes covering 38,256,468 unique SNPs. These phenotypes serve as the final analytical units for PRS calculation, risk stratification, and individual-level interpretation. The generated reports integrate sample-level results, phenotype categorization, risk classification, study references, and variant tables, providing a structured and interpretable output for end users. Together, the curated database and reporting framework establish a comprehensive toolbox for PRS analysis, enhancing reproducibility, transparency, and usability in both research and clinical contexts.

## Introduction

The rapid decrease in the cost of DNA sequencing over the past two decades has revolutionized biomedical research and clinical practice. What was once prohibitively expensive is now widely accessible, enabling the generation of vast amounts of genomic data from both healthy individuals and patients. This technological advance has fueled the development of numerous genetic tests, which are currently applied to a broad spectrum of normal physiological processes as well as pathological conditions. The primary goal of these tests is to identify genetic alterations, whether germline variants inherited across generations or somatic mutations acquired during life, that can improve our understanding of metabolism, disease mechanisms, and therapeutic responses. Among the most significant outcomes of this genomic revolution is the emergence of precision medicine, a paradigm that tailors diagnosis, prevention, and treatment strategies to the unique genetic makeup of each individual (1).

Within this context, one particularly promising approach for predicting phenotypic features from genetic data is the calculation of polygenic risk scores (PRS) (2, 3). Unlike single-gene tests that focus on rare variants with large effects, PRS aggregate the influence of many common genetic variants across the genome. Each variant contributes a small effect size, but collectively they can provide meaningful insights into an individual’s predisposition to complex traits and diseases (4, 5). PRS are calculated by comparing an individual’s genotype to sets of polymorphisms statistically associated with a trait, as identified through genome-wide association studies (GWAS). In this framework, risk is interpreted as a relative position within a reference population rather than an absolute prediction of disease occurrence.

The calculation of PRS typically relies on whole-genome sequencing (WGS), which provides comprehensive coverage of genetic variation. However, whole-exome sequencing (WES) can also be used when complemented by genotype imputation to infer variants outside captured exonic regions. Advances in statistical methods and computational tools have further refined PRS estimation, improving accuracy and portability across different datasets. Despite these improvements, challenges remain in ensuring that PRS are robust across diverse populations, given the historical predominance of European-ancestry cohorts in GWAS and score development (6, 7).

To support the development and application of PRS, several curated databases and computational pipelines have been established. The NHGRI-EBI GWAS Catalog provides a comprehensive repository of published associations between genetic variants and traits (8), whereas the Polygenic Score (PGS) Catalog systematically reports score models and their associated metadata (9). Tools such as PRSice-2 and STREAM-PRS have also contributed to the standardization and automation of PRS workflows (10, 11). Here, we present a pipeline designed to integrate data from the GWAS Catalog and the PGS Catalog, calculate PRS for multiple traits, estimate relative risk by comparison with population-level distributions, and produce structured individual reports.

## Methods

### Pipeline overview

We developed an end-to-end computational pipeline for the construction of a curated genotype/phenotype database, calculation of polygenic risk scores in individual samples, population-based risk stratification, and automated report generation. The workflow integrates associations from GWAS, published polygenic score models, genotypes from a reference population, and variants observed in the analyzed individual. The pipeline was divided into two complementary stages: database construction and individual-level execution (Figure 1).

**Fig 1.**
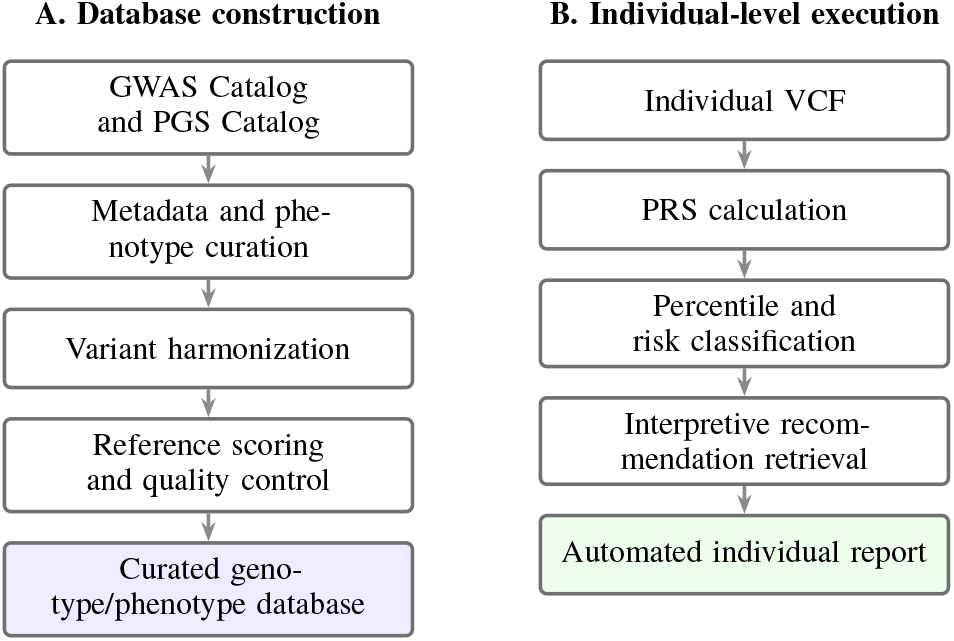
Overview of the computational pipeline. Database construction and individual-level execution are performed as complementary stages.

During database construction, GWAS associations and published score models are curated, harmonized, and linked to phenotype and study metadata. Variants are standardized to a common genomic representation, and the resulting phenotype-specific models are applied to the 1000 Genomes Project to construct reference distributions. During individual-level execution, an input VCF is queried against the curated database, PRS are calculated, and each score is positioned within the corresponding global and population-stratified distributions. The resulting percentiles, risk classes, study information, detected variants, and previously generated interpretive texts are then assembled into an automated report.

Separating database construction from individual analysis avoids repeating computationally intensive or potentially variable procedures for every new sample. Reference distributions and language-model outputs are therefore generated once and reused during subsequent analyses.

### Data sources and construction of the curated database

The genotype/phenotype database was constructed using the GWAS Catalog and the PGS Catalog, which provide complementary information for PRS analysis (8, 9). The GWAS Catalog was used as a source of published variant–trait associations, whereas the PGS Catalog provided published score models, including effect alleles, statistical weights, and model-development metadata.

Because the source files use heterogeneous structures, records were converted to a common tabular format. Each variant was represented by the genomic key `CHR:POS:REF:ALT`and linked to the effect allele, statistical weight, rsID, and gene or genomic context when available. This coordinate- and allele-based representation allowed direct comparison with VCF files and reduced dependence on rsIDs that may be absent or inconsistent between reference versions.

VCF manipulation and genomic harmonization were performed with *bcftools* (12), and functional context was annotated with *snpEff* (13). Harmonization included removal of the `chr`prefix, coordinate compatibility checks, and confirmation of reference and alternate alleles. Native PGS Catalog identifiers were preserved, whereas GWAS-derived entries received internal identifiers linking each phenotype to its variants, study metadata, reference distributions, and report-related information.

The curation process included phenotype-name standardization, consolidation of descriptions and bibliographic information, thematic categorization, identification of applicability by biological sex, removal of duplicates, and exclusion of phenotypes considered unsuitable for direct presentation in an individual report. These exclusions included traits with limited interpretability, sensitive content, or inadequate suitability for user-facing communication. When multiple entries represented the same phenotype, preference was given to the entry containing the largest number of associated variants. A language model supported semantic normalization and preparation of interpretive metadata, with temperature set to zero to reduce variability. Final inclusion also depended on structural and statistical quality-control rules implemented in the pipeline.

### Reference population and distribution quality controll

The reference population was obtained from the 1000 Genomes Project, which comprises genetic variation from multiple populations and superpopulations (14). For each curated phenotype, the corresponding variants, effect alleles, and weights were applied to the reference genotypes with PLINK2 (15). This procedure generated one score per reference individual for each phenotype.

Reference distributions were constructed for the complete cohort and separately for the AFR, AMR, EAS, EUR, and SAS superpopulations. The mean (*µ*) and standard deviation (*σ*) were calculated for each phenotype and population group, allowing the same individual PRS to be interpreted relative to the complete cohort and to each major superpopulation represented in the dataset.

Before inclusion in the curated database, population score distributions were evaluated with the Shapiro–Wilk test (16, 17). Under the operational criterion adopted in the pipeline, distributions with a *p*-value below 0.93 were excluded. This step restricted the subsequent parametric calculation of z-scores and percentiles to distributions considered compatible with the use of the mean, standard deviation, and cumulative standard normal distribution.

### Individual sample processing and PRS calculation

The pipeline uses a VCF containing the genotypes of the analyzed individual. WGS data can be processed directly. WES data are processed after genotype imputation with Beagle using the 1000 Genomes Project as a reference panel (18). This step expands the number of evaluable loci outside captured coding regions, although imputed genotypes remain subject to uncertainty.

Individual variants were standardized with the same `CHR:POS:REF:ALT`representation used in the curated database. For each phenotype, the pipeline intersected the associated variants with the evaluable variants in the individual sample. The resulting records included the genomic key, observed genotype, effect allele, statistical weight, rsID, and associated gene.

Genotypes were converted into effect-allele dosages. A genotype without the effect allele received a dosage of 0, a heterozygous genotype received a dosage of 1, and a genotype homozygous for the effect allele received a dosage of 2. Equation 1 presents the normalized PRS formulation used in this work.

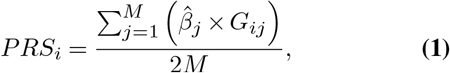

where *M* is the number of phenotype-associated variants considered in the calculation, 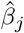 is the estimated effect of variant *j*, and *G*_*ij*_ is the effect-allele dosage for individual *i* (0, 1, or 2). The denominator 2*M* corresponds to the maximum number of alleles evaluated for the *M* variants in a diploid organism. Applying the same normalization rule to the individual sample and the reference population maintains a consistent scale for population-based positioning. Figure 2 illustrates the calculation using three phenotype-associated variants.

**Fig 2.**
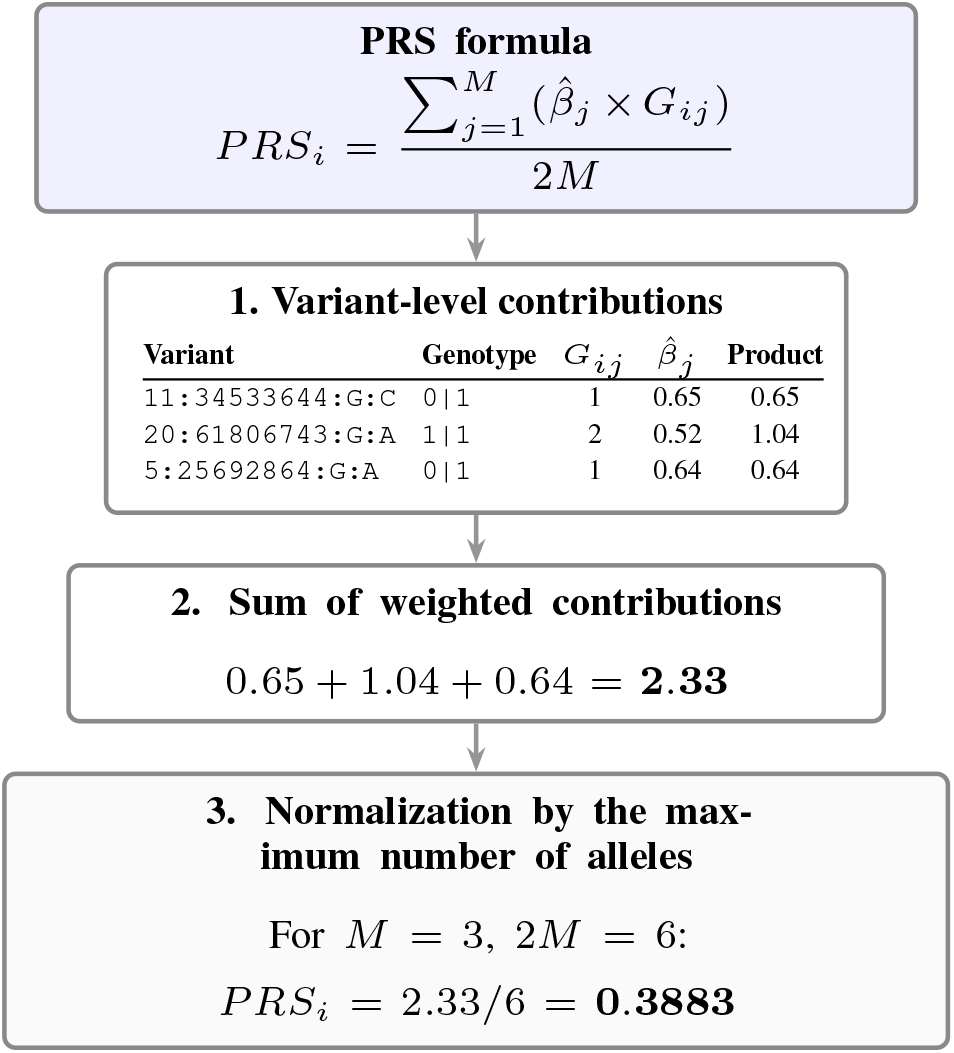
Step-by-step PRS calculation. Effect-allele dosages are multiplied by their statistical weights, summed, and normalized by the maximum number of evaluated alleles.

Phenotypes without detected effect variants in the individual were not included in the final report, preventing risk classifications based on an empty marker set.

### Percentile estimation and risk classification

For each phenotype, the individual PRS was compared with the corresponding reference distribution. The standardized score was calculated as

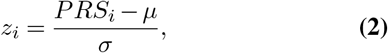

where *µ* and *σ* are the mean and standard deviation of the reference distribution. The z-score was then converted into a percentile using the cumulative distribution function of the standard normal distribution:

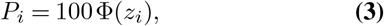

where Φ(*z*_*i*_) is the cumulative proportion located to the left of the individual score. The calculation was performed for the complete cohort and separately for each available superpopulation.

Percentiles were divided into five relative risk levels (Table 1). These categories describe the relative position of an individual’s score and do not represent the absolute probability of developing a disease or trait.

**Table 1.** Percentile intervals used for relative risk classification.

| Percentile interval | Risk level |
| --- | --- |
| $P < 10$ | Very low |
| $10 \leq P < 25$ | Low |
| $25 \leq P < 75$ | Medium |
| $75 \leq P < 90$ | High |
| $P \geq 90$ | Very high |

### Generation of interpretive recommendations

Interpretive recommendations were generated in advance with a large language model rather than during sample processing. For each phenotype, the normalized name, description, study summary, identifier, and one of the five risk levels were provided to GPT-4o-mini through an API. The model was configured with temperature set to zero, and LangFlow was used only to orchestrate the generation workflow.

The generation criteria required the model to account for the direction and meaning of each phenotype, avoid assuming that a high value was necessarily unfavorable or that a low value was necessarily protective, use accessible language, and avoid diagnoses, prescriptions, or deterministic statements. Five texts were generated for each phenotype and stored in a JSON structure indexed by phenotype identifier and risk level. During individual analysis, the pipeline retrieves the corresponding pre-generated text without issuing a new request to the model. These texts support interpretation and communication but do not replace medical evaluation or genetic counseling.

### Automated report generation and computational implementation

Results for each individual were consolidated in a JSON structure containing sample information, PRS values, z-scores, global and stratified percentiles, risk levels, phenotype metadata, recommendations, and detected variants. The report was organized by thematic category and risk level and included a sample overview, phenotype summaries, individual phenotype pages, study metadata, population-specific comparisons, interpretive recommendations, and tables of detected variants.

The implementation was divided into modules for database preparation, variant processing, score calculation, statistical classification, recommendation retrieval, and report generation. Python and shell scripts were used for tabular processing and workflow control, Crystal was used for high-performance variant intersection, *bcftools* and Beagle handled genomic preprocessing, and PLINK2 generated reference-population scores. Intermediate and final data were exchanged through CSV, TSV, VCF, and JSON files. The modular structure allows database preparation to be performed independently of individual analysis and permits phenotype-level tasks to be parallelized. Our pipeline is available for non-commercial users upon request.

## Results

### Construction of the Curated Phenotype Database

The first stage of the results corresponds to the construction of the curated genotype/phenotype database employed by the PRS pipeline. As outlined in the methodology, data obtained from public repositories were initially standardized, organized by unique identifiers, and subjected to a curation process prior to individual-level analysis. This curation aimed to ensure that only phenotypes with appropriate structure, interpretable metadata, and compatible population distributions were retained in the final dataset.

In total, 13,423 studies associated with phenotypes were processed. For each, polygenic scores were calculated in the reference population, and the distribution of population scores was assessed. This step was critical, since the pipeline relies on the relative position of an individual within the reference population to compute z-scores, percentiles, and risk levels. Among the processed studies, 5,543 exhibited distributions classified as normal according to the pipeline’s criteria and were retained in the curated database. Conversely, 7,880 displayed non-normal distributions and were excluded. This filtering represented a central stage of the curation process, as it eliminated phenotypes whose population scores did not conform to the inference strategy based on mean, standard deviation, z-score, and percentile.

Following the evaluation of population score distributions, the remaining phenotypes underwent further curation steps. These included the removal of duplicates, standardization of metadata, normalization of phenotype names, thematic categorization, and exclusion of phenotypes deemed unsuitable for presentation in a user-facing report. This stage was necessary because not all phenotypes available in public repositories are appropriate for inclusion in an individual interpretive report. Certain phenotypes may represent sensitive traits, provide limited information, be difficult to interpret, or be ethically inadequate for direct communication to end users.

At the conclusion of this curation process, the consolidated database used by the pipeline contained 2,346 phenotypes. These phenotypes represent the final analytical units available for comparison with individual data, PRS calculation, risk classification, and report generation. Moreover, the final database encompassed 38,256,468 unique SNPs, reflecting the total set of variants associated with the retained phenotypes. This breadth of coverage defines the scope of variants that can be queried in individual samples during the intersection between patient VCF files and the curated phenotype variant files.

**Table 2.**
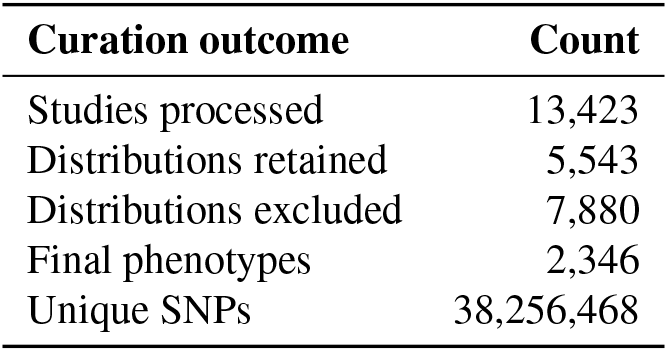
Summary of the curated genotype/phenotype database.

### Automated Generation of Individual PRS Reports

In addition to the construction of the curated genotype/phenotype database, another central outcome of this work was the automated generation of individual polygenic risk score reports. This report represents the final output of the pipeline, integrating the results calculated for each individual, the organization of phenotypes by thematic category, the estimated risk levels, associated recommendations, source study information, and the genetic variants detected in the sample.

To illustrate the structure of the generated report, an example document produced by the pipeline was used. The selected components highlight: i) sample overview – general information about the analyzed individual; ii) categorical organization – grouping of phenotypes into thematic categories; iii) risk-level listing – phenotypes ordered according to estimated risk levels; iv) individual phenotype pages – detailed information for each phenotype, including risk classification and study metadata; and v) variant tables – genetic variants associated with the retained phenotypes. Figure 3 presents a representative phenotype page, including the population-specific comparison and the interpretive recommendation generated with the language model, followed by the associated variant table.

**Fig 3.**
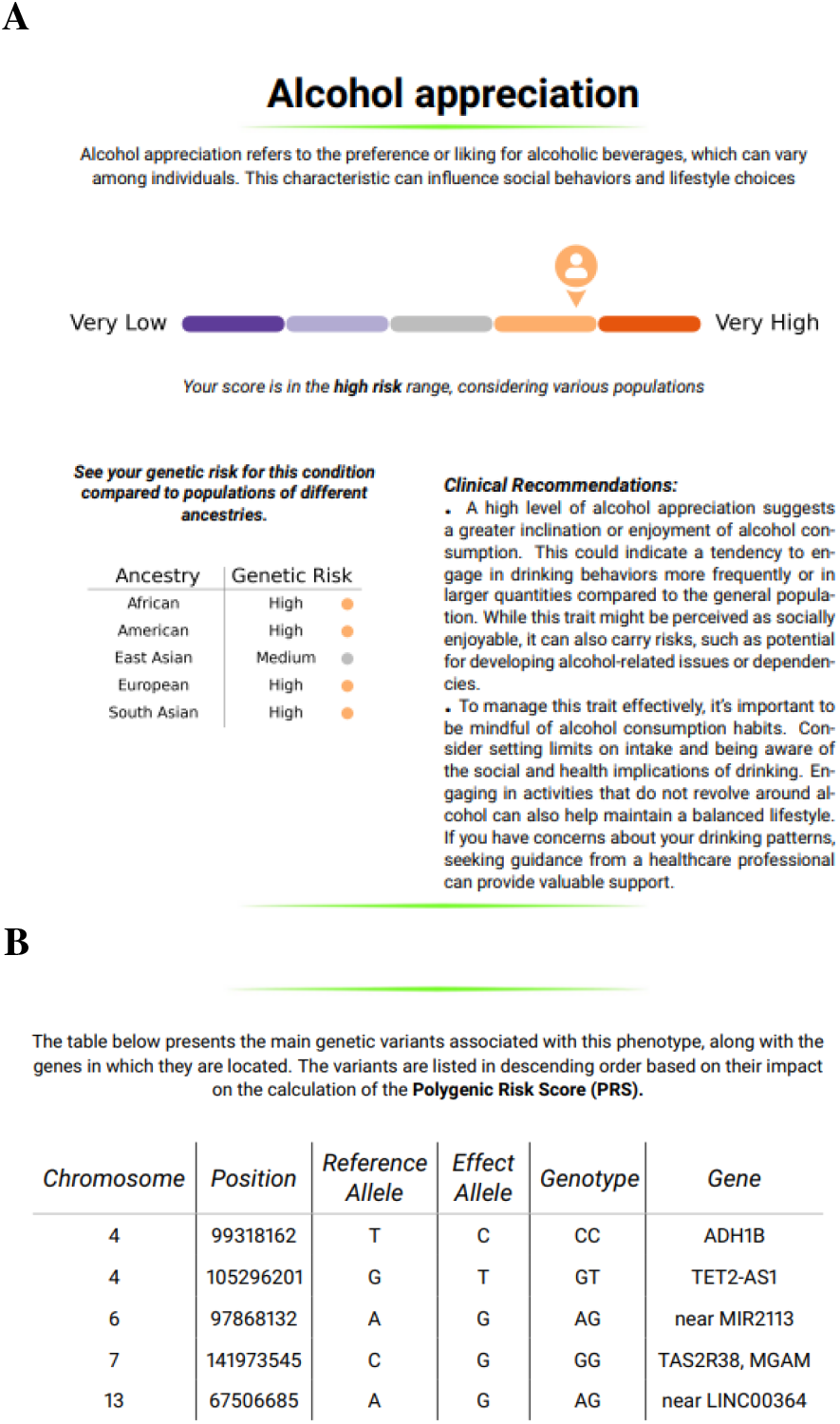
Representative outputs of the automated individual PRS report: (A) a phenotype-specific page containing the percentile-based risk scale, population-specific comparison, and LLM-generated interpretive recommendation; and (B) the table of detected variants contributing to the phenotype-specific result.

## Discussion

In this work, we present a pipeline designed to generate polygenic risk scores for traits available in the GWAS Catalog and PGS Catalog and to estimate individual risk through population stratification and comparative analysis. The resulting report provides, for each trait, the list of SNPs contributing to the PRS, the calculated PRS value, and the corresponding risk assessment derived from comparisons with the distribution of PRS in the 1000 Genomes Project cohort. A distinctive feature of our approach is that, in addition to evaluating risk relative to the entire 1000 Genomes population, the pipeline also enables stratified analyses across subsets of the cohort defined by major population groups. This functionality enhances the interpretability of PRS by allowing the identification of individuals whose genetic risk is particularly pronounced within the context of their population background.

Several methodological considerations must be acknowledged. A primary concern is the reliance on imputation for datasets derived from WES. Although imputation has been successfully applied across diverse genomic contexts, inaccuracies in imputed genotypes can compromise PRS estimation, particularly when errors involve SNPs with large effect sizes (19). Another important limitation relates to the stratification of population groups in admixed individuals. In our analyses, we adopted the framework used by the 1000 Genomes Project, in which geographic origin is a strong component. This approach, although widely employed, may oversimplify the complex genetic structure of admixed populations and potentially obscure more nuanced ancestry-related risk patterns (6, 7).

Despite these limitations, our pipeline provides several significant advantages, particularly in its flexibility for selecting traits to be evaluated and its emphasis on methodological standardization. It produces comprehensive and informative reports that not only summarize key findings but also present them in a clear, well-structured, and user-friendly format. The separation between database construction and individual execution also promotes reproducibility by ensuring that reference distributions and interpretive texts are prepared once and reused under the same operational rules. Standardized reporting remains an important component for the transparent interpretation and communication of polygenic scores (20).

